# Multi-organ Metabolic Model of *Zea mays* Connects Temperature Stress with Thermodynamics-Reducing Power-Energy Generation Axis

**DOI:** 10.1101/2023.07.09.548275

**Authors:** Niaz Bahar Chowdhury, Berengere Decouard, Isabelle Quillere, Martine Rigault, Karuna Anna Sajeevan, Bibek Acharya, Ratul Chowdhury, Bertrand Hirel, Alia Dellagi, Costas Maranas, Rajib Saha

## Abstract

Global climate change has severely impacted maize productivity. A holistic understanding of metabolic crosstalk among its organs is essential to address this issue. Thus, we reconstructed the first multi-organ maize genome-scale metabolic model, *i*ZMA6517, and contextualized it with heat and cold stress-related transcriptomics data using the novel **<u>EX</u>**pression dis**<u>T</u>**ributed **<u>REA</u>**ction flux **<u>M</u>**easurement (EXTREAM) algorithm. Furthermore, implementing metabolic bottleneck analysis on contextualized models revealed fundamental differences between these stresses. While both stresses had reducing power bottlenecks, heat stress had additional energy generation bottlenecks. To tie these signatures, we performed thermodynamic driving force analysis, revealing thermodynamics-reducing power-energy generation axis dictating the nature of temperature stress responses. Thus, for global food security, a temperature-tolerant maize ideotype can be engineered by leveraging the proposed thermodynamics-reducing power-energy generation axis. We experimentally inoculated maize root with a beneficial mycorrhizal fungus, *Rhizophagus irregularis*, and as a proof of concept demonstrated its potential to alleviate temperature stress. In summary, this study will guide the engineering effort of temperature stress-tolerant maize ideotypes.

---

Temperature stress, resulting from global climate change, can reduce maize productivity by 7-18%^1^. Thus, there is a pressing need to develop high-yielding maize genotypes capable of withstanding temperature stress. A metabolism-centric approach can be useful to achieve that. Thus, an in-depth understanding of the impact of temperature on plant-wide metabolism is required. Such impacts include, but are not limited to, reduced photosynthesis and carbohydrate synthesis in leaves^2^, reduced starch synthesis in kernels^3^, upregulation of amino acid and downregulation of diterpenoid metabolism in root^4^, and lignin biosynthesis in stalks^5^. Although these studies were informative, a holistic plant-wide understanding of temperature stress responses, delineating the interactions between vegetative and reproductive organs at key stages of plant development, is still in its infancy. Moreover, the plant-wide effect of well-known beneficial arbuscular mycorrhizal fungi (AMF), such as *Rhizophagus irregularis*^6^, has not been evaluated for its potential to alleviate temperature stress on maize growth. A multi-organ genome-scale metabolic model (GSM) is suited to address these issues. The first multi-organ plant GSM was reconstructed for barley^7^, subsequently for barrelclover^8^, arabidopsis^9,10^, soybean^11^, foxtail millet^12^, and rice^13^. These models were useful to characterize inter-organs crosstalk under various conditions. Thus, for a holistic understanding of plant-wide temperature stress responses of maize, we reconstructed the first multi-organ maize GSM, *i*ZMA6517, integrating comprehensive root, stalk, kernel, and leaf GSMs.

Contextualization of GSM by integrating ‘omics’ data is a crucial step to connect metabolism to phenotypes. There are two classes of algorithms for contextualization: valve approaches (assuming reactions fluxes and transcript levels are proportional) and switch approaches (assuming reaction are turned on/off based on transcript levels). Earlier studies discussed the suitability of both approaches^14^, showing E-flux algorithm^15^ (a valve approach) capturing better phenotypic predictions. However, E-flux algorithm can predict unrealistic phenotypes too^16^. Thereby, to improve the phenotypic prediction accuracy, there is a need to develop a novel valve approach which can address this limiation. To achieve this, we proposed the EXTREAM algorithm. Moreover, we propose a Metabolic Bottleneck Analysis (MBA) algorithm to pinpoint metabolic bottlenecks in *i*ZMA6517.

In this study, we combined *i*ZMA6517, EXTREAM, and MBA to dissect plant-wide responses of temperature stress by pinpointing metabolic bottlenecks in different organs and to determine if responses to cold and heat share common characteristics. We found reducing power capacity had a pivotal role in responses to both stresses. Additionally, heat stress responses incurred energy generation bottlenecks. Thermodynamic driving force analysis of bottleneck pathways highlighted the role of thermodynamics-reducing power-energy generation axis when plants are subjected to temperature stress. Finally, as a proof of concept, we extended our analysis by integrating maize “omics” data when the plant is inoculated with the AMF *R. irregularis* in *i*ZMA6517 and found that the inoculation can alleviate temperature stresses. Ultimately, this study could offer a blueprint to engineer robust abiotic stress-tolerant maize ideotypes.

## RESULTS

### Maize responses to heat and cold stress

To decipher maize responses to temperature stress, heat and cold-stress-related transcriptomic data of B73 genotype were collected from the literature^17^. Seedlings were subjected to cold (5°C for 16 hours) or heat (50°C for 4 hours) stresses, compared to a control condition (24°C). All plants were grown in the autoclaved field soil. While heat stress drastically affected transcriptional response compared to control conditions (Fig. 1A), the effect of cold stress was similar to control conditions (Fig. 1B), indicating a difference of plant adaptations to temperature stresses (Fig. 1C, D). To explore the components of temperature stresses, we performed K-means clustering (Fig. 1E) on both heat and cold stress data, identifying four distinct clusters followed by gene enrichment analyses (Supplementary Fig. 1-4). These analyses revealed that photosynthesis-related genes (cluster 2) were upregulated under both stress conditions. In addition, heat shock genes (cluster 3) were upregulated in heat stress conditions. Cluster 1 and 2 revealed genes associated with central carbon and secondary metabolism, respectively.

**Fig. 1.**
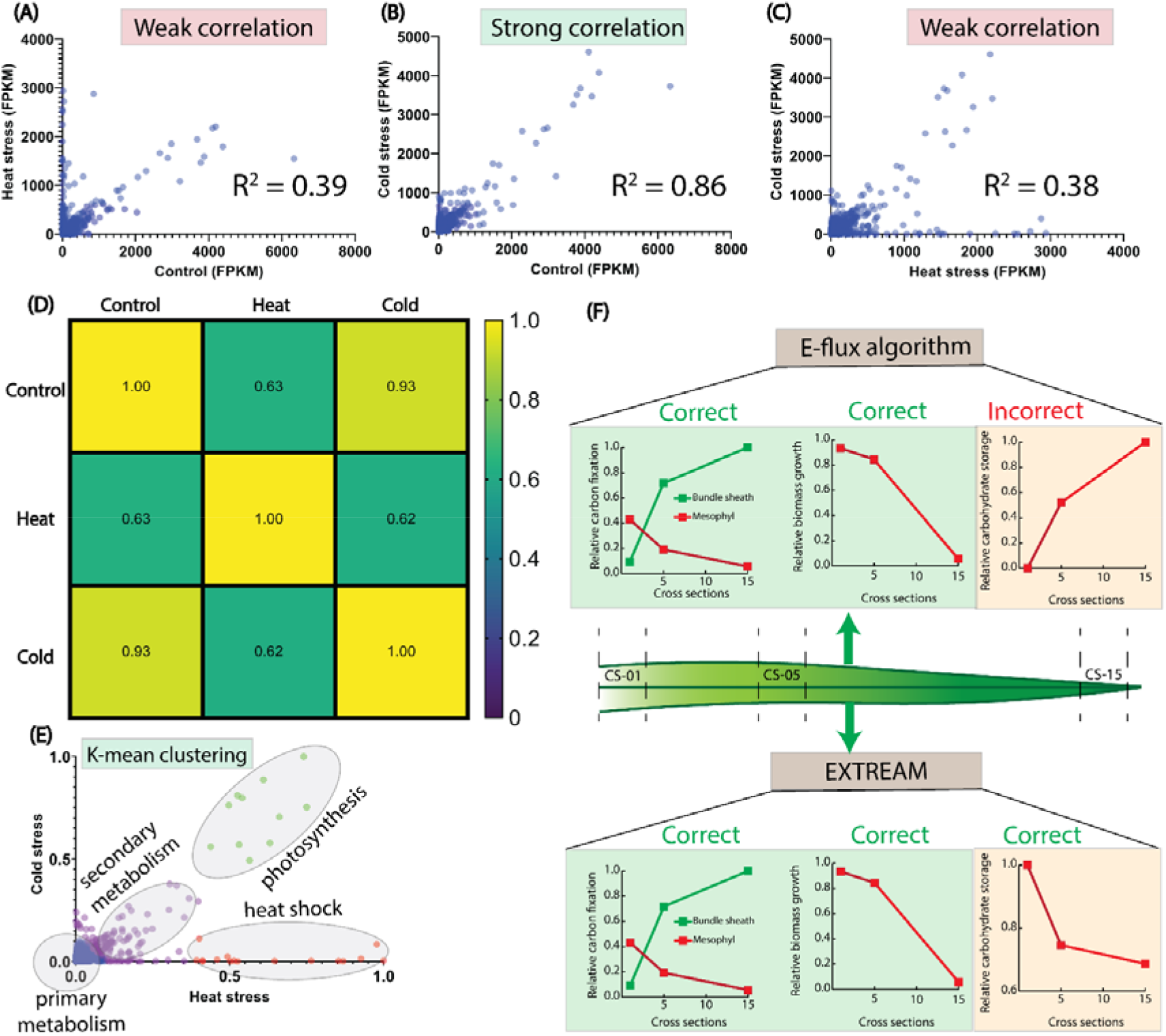
Transcriptomics data analysis and introduction to EXTREAM algorithm. A) Scatter plot of control and cold stress transcriptomics data. B) Scatter plot of control and heat stress transcriptomics data. C) Scatter plot of heat and cold stresses transcriptomics data. D) Correlation matrix among control, heat stress, and cold stress (95% confidence interval, two-tail test). E) K-means clustering analysis of heat and cold stress data. F) E-flux and EXTREAM both predicted correct biomass production and carbon fixation for three selected cross sections of leaf, whereas only EXTREAM predicted the correct leaf starch content.

As photosynthesis is one of the core metabolic features of plants, cluster 2 was examined in further detail. A previous study indicated a similar high transcriptional response of photosynthetic genes under temperature stress^18^. However, photosynthetic activity of plants, as measured through decreased rubisco activation^19^, decreases under temperature stress. This suggests potential metabolic bottlenecks in plant tissues preventing higher photosynthetic activity in leaves. A contextualized multi-organ GSM of maize can identify such bottlenecks. There may also be bottlenecks between transcription and translation, such as mRNA degradation, however, that would be outside of the scope of GSM. Next, we will design a suitable algorithm for contextualization of GSM.

### EXTREAM algorithm

Our previous study used the E-flux algorithm^20^ to accurately predict the maize root phenotype under nitrogen starvation based on transcriptomic data. This work involved photosynthetic tissues, having additional metabolic complexities compared to roots. Thereby, the efficacy of the E-flux algorithm was re-assessed.

In a previous study, the maize leaf was divided into 15 cross-sections, and generated transcriptomics data for each of these sections^21^. For this study, we collected transcriptomics data of cross-sections 1, 5, and 15 based on the greenness of the leaf blade (less green to greener). These data were integrated in the leaf GSM^22^ using E-flux algorithm. The highest growth rates in maize leaves occur at the base of the leaf, where new cells are actively dividing and elongating. As the cells grow toward the tip, they undergo expansion and maturation, leading to a decrease in the growth rate. Eventually, the cells at the tip stop elongating altogether. The contextualized leaf model correctly predicted higher biomass growth rate of leaf base compared to leaf tip (Fig. 1F). The model also predicted rate of carbon fixation correctly, however, it incorrectly predicted the starch distribution profile along the leaf (Fig. 1F). Thus, modification of the E-flux algorithm was required to fit the starch distribution profile.

To achieve this, we proposed the EXTREAM algorithm, which equally distributed transcript levels for each gene based on the number of cognate biochemical reactions. Once we contextualized the leaf model with the EXTREAM algorithm, it predicted the correct biomass growth rate, carbon fixation, and starch distribution profiles along the selected leaf sections (Fig. 1F). Thus, EXTREAM algorithm can be effective in modeling plant organs, including photosynthetic organs.

### Multi-organ genome-scale metabolic model of maize

In this study, we reconstructed the first multi-organ maize GSM, *i*ZMA6517 (Fig. 2A) for the B73 genotype. The model is based on the previously reconstructed root and leaf-specific GSMs^14,22^, and newly reconstructed stalk and kernel GSMs, connected *via* vascular tissues (Supplementary Data 1).

**Fig. 2.**
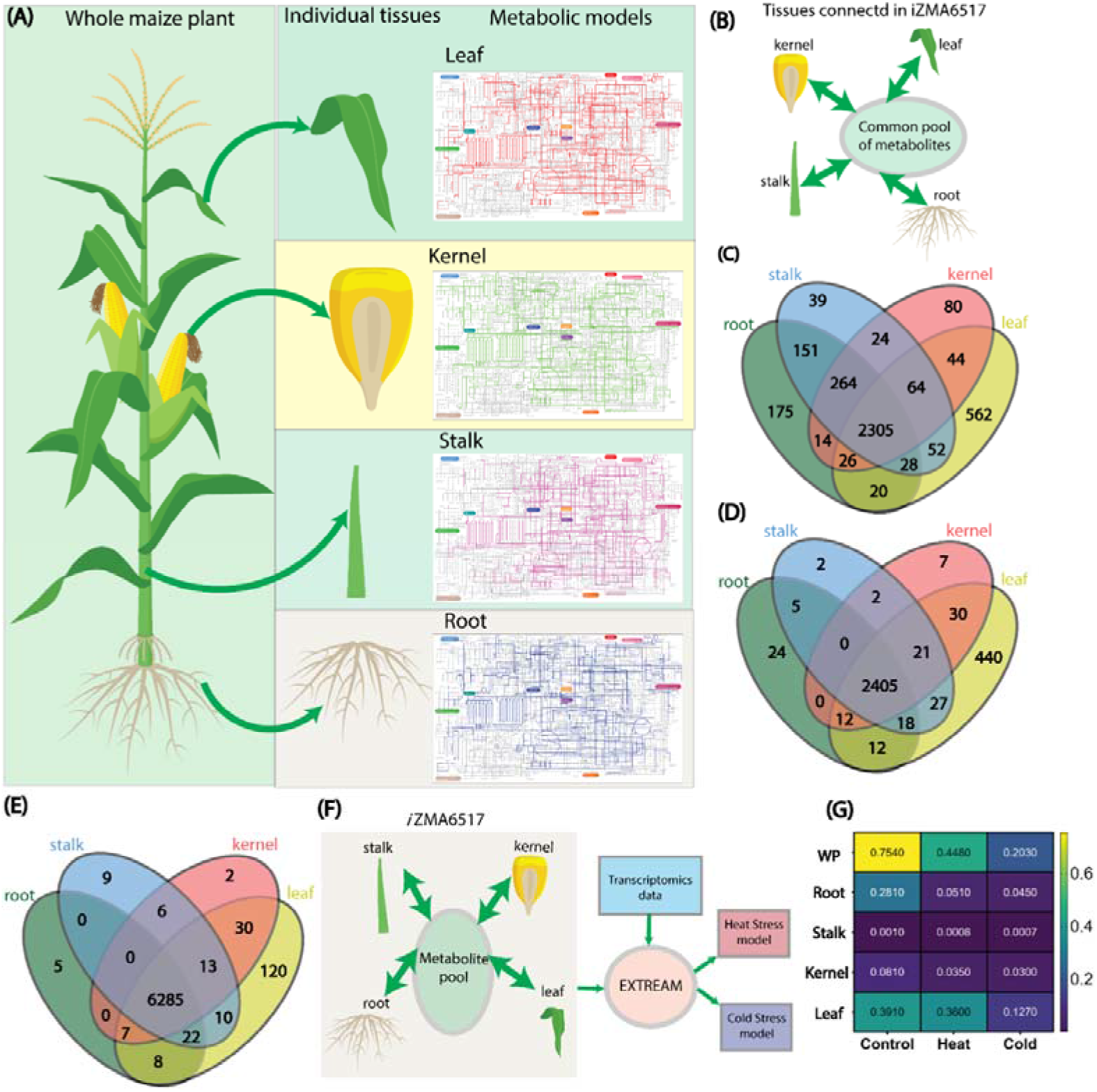
Overview of *i*ZMA6517 reconstruction and contextualization. A) Individual metabolic models for root, stalk, kernel, and leaf. B) Connection between individual tissues via the vascular tissues. C) Comparison of reactions among the individual GSMs. D) Comparison of metabolites among the individual GSMs. E) Comparison of unique genes among the individual GSMs. F) Incorporating heat and cold stress related transcriptomics data with the iZMA6517 using the EXTREAM algorithm. G) EXTREAM algorithm predicted lower biomass production in each organ and in the whole plant (WP) during the cold stress compared to the heat stress.

To further refine our existing leaf model, we identified thermodynamically infeasible cycles (TICs) and resolved those using our previously-developed OptFill^23^ pipeline. For stalk and kernel GSMs, we assigned gene-protein-reaction (GPR) relationships to all known reactions based on the MaizeCyc databse^24^. The GPR relationships used Boolean logic to determine if enzymes associated with metabolic reactions are encoded by isozymes (OR relationship) or protein complexes (AND relationship). We next identified organ-specific reactions and the corresponding metabolic pathways by categorizing each gene’s expression level for each organ. Next, we reconstructed the stalk and kernel biomass equation based on literature (Supplementary Data 2). Upon completing the pathway participation in the organ-specific GSMs, we carried out gap-filling (Supplementary Note).

Finally, *i*ZMA6517 was assembled by connecting organ-specific GSMs *via* vascular tissues, namely phloem and xylem (Fig. 2B). Although phloem and xylem are distinct tissues, the direction of the flow of metabolites was sufficient to determine if it was present in either one of those. Therefore, the two tissues were combined for simplicity. These vascular tissues will facilitate inter-organ crosstalk by translocating metabolites from one organ to another. For example, sugars travel from the leaf tissue to other tissues *via* the vascular tissues. Similarly, nutrients, such as nitrate and phosphate, are uptaken by root and distributed to the other organs. Supplementary Data 3 has details on inter-organ transfer of metabolites. Overall, *i*ZMA6517 contains 6517 genes, 5228 unique reactions, and 3007 unique metabolites. Among these, 2305 reactions (Fig. 2C), 2405 metabolites (Fig. 2D), and 6285 genes (Fig. 2E) were common across all organs.

Next, to assess plant-wide impact of temperature stress, we contextualized *i*ZMA6517 for heat and cold stresses using the EXTREAM algorithm (Fig. 2F). The transcriptomics data from the seedling were the aggregate data of root, stalk, and leaf. As kernel is a sink, like a previous study where it showed that there is a control of assimilates translocation from stalk to kernels^25^, we assumed the metabolism of kernel will be dictated by the aggregate behavior of root, stalk, and leaf. Thus, the aggregate transcriptomics data were used to contextualize the model (*i*ZMA6517) comprising of root, stalk, kernel, and leaf under heat and cold stress. Previous studies showed control conditions had the highest nutrient uptakes, followed by heat and cold stress^26^. Once these nutrient uptakes patterns were implemented in contextualized *i*ZMA6517s, control and cold-stressed plants showed the highest and lowest biomass growth rates, respectively (both individual organs and whole plant) (Fig. 2G), supporting previous studies^27^. Indeed, by projecting the aggregate transcriptomics data from seedling to kernel, we were able to predict the correct growth rate pattern for kernel in control, heat stress, and cold stress (Fig. 2G). That confirms our initial assumption that, as a sink, the kernel metabolism is dictated by other organs like root, stalk, and leaf. Thus, the contextualized *i*ZMA6517 were suitable for identifying metabolic bottlenecks of temperature stresses.

### Identification of metabolic bottlenecks

To assess the impact of temperature stress on maize metabolism, we implemented Metabolic Bottleneck Analysis (MBA) to contextualized *i*ZMA6517s. MBA expanded the flux space of each reaction separately and assessed its impact on the whole plant biomass growth rate. It revealed 180 bottleneck reactions under heat stress, of which 70% occurred in leaves, 28% in kernels, and 2% in roots (Fig. 3A). These reactions were distributed across purine metabolism, pyrimidine metabolism, fatty acid metabolism, the Calvin cycle, and glycolysis (Supplementary Fig. 5). Among these 180 reactions, root cytochrome b5 reductase (24%) and acyl-ACP-hydrolase (24%) increased the plant biomass growth rate the most. Here, percentages indicate increase in biomass growth rate after debottlenecking a specific bottleneck reaction, compared to the biomass growth rate of corresponding stress condition before debottlenecking. Overall, 19 reactions increased plant biomass growth rate by more than 10% (Supplementary Fig. 6).

**Fig. 3.**
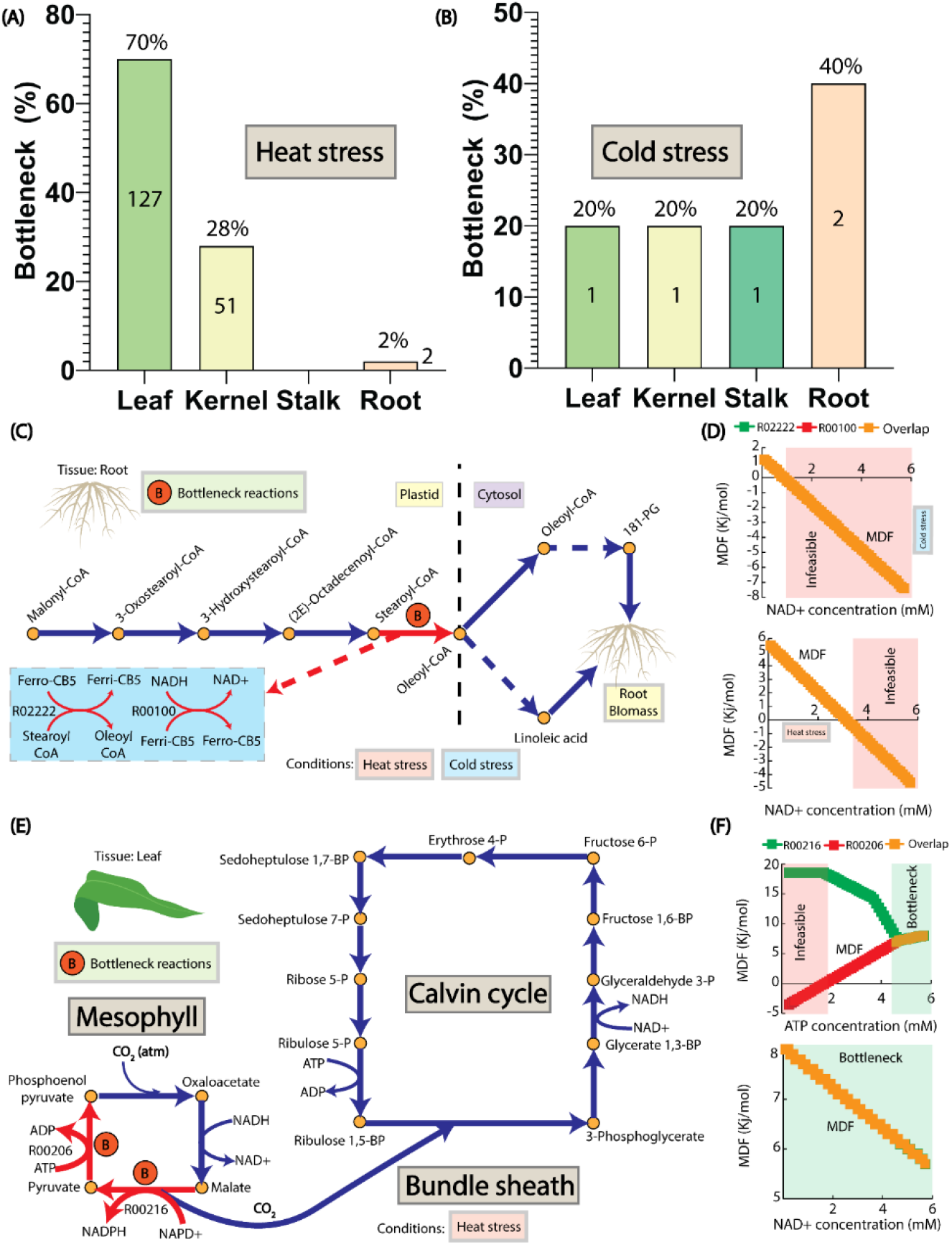
Identification of temperature stress bottlenecks. A) Organ specific bottleneck reactions for heat stress (number of reactions is indicated inside the column). B) Organ specific bottleneck reactions for cold stress, (number of reactions is indicated inside the column). C) Cytochrome b5 reductase bottleneck in the fatty acid metabolism for heat and cold stress. D) Thermodynamic driving force analysis in fatty acid metabolism. E) Pyruvate-phosphate dikinase and malate dehydrogenase bottlenecks in the photosynthetic pathway for heat stress. F) Thermodynamic driving force analysis of the photosynthetic pathway.

For cold stress, MBA revealed five different bottleneck reactions Two of these occurred in the root, and one each in other organs (Fig. 3B). Among those five reactions, similar to the heat stress, cytochrome b5 reductase in roots (182%) and leaves (147%) and acyl-ACP-hydrolase in roots (146%) increased the plant biomass growth rate the most. The stalk coniferyl-aldehyde dehydrogenase (32%) and the kernel phosphohexomutase (4%) were the other bottleneck reactions.

As reaction thermodynamics is influenced by temperature, we applied Min/Max Driving Force (MDF) analysis to pathways containing bottleneck reactions, within the physiological range of substrate concentration (0.01 mM to 10 mM)^28^ to assess the impact of thermodynamics on those pathways (Supplementary Note). The standard Gibbs free energy of reactions was calculated (Supplementary Data 4) using the group contribution method^29^. The root cytochrome b5 reductase and acyl-ACP-hydrolase were common to both temperature stresses. The reaction catalysed by cytochrome b5 reductase is part of the fatty acid biosynthesis pathway (Fig. 3C) and impacted the production of 18:1-phosphoglycerate choline and linoleic acid of biomass components. MDF analysis indicated that, with an NAD^+^ concentration greater than 1.1 mM (cold stress) and 3.7 mM (heat stress), the fatty acid biosynthesis pathway was thermodynamically feasible and the cytochrome b5 reductase had the lowest driving force (Fig. 3D). Acyl-ACP-hydrolase is involved in the fatty acid biosynthesis pathway and impacted the octadecanoic acid of biomass components (Supplementary Fig. 7). MDF analysis again revealed that the pathway was thermodynamically feasible when the NAD^+^ concentration was between 0.1 to 5.7 mM, while acyl-ACP-hydrolase had the lowest driving force (Supplementary Fig. 8).

Coniferyl-aldehyde dehydrogenase involved in the phenylpropanoid biosynthesis pathway (Supplementary Fig. 9) in stalk was a cold stress-related bottleneck and impacted the production of ferulic acid of the stalk biomass. MDF analysis revealed that, with an NAD^+^ concentration greater than 0.3 mM, the phenylpropanoid biosynthesis pathway became thermodynamically infeasible, and coniferyl-aldehyde dehydrogenase had the lowest driving force (Supplementary Fig. 10). Phosphohexomutase of the fructose-mannose biosynthesis pathway in the kernel was a cold stress bottleneck (Supplementary Fig. 11). MDF analysis showed that the fructose-mannose biosynthesis pathway was thermodynamically feasible only when the concentration of fructose 6-phosphate was between 2.5 and 5 mM. In this concentration range, phosphohexomutase was the reaction with the lowest driving force (Supplementary Fig. 12).

For heat stress-only metabolic bottlenecks, we focused on pyruvate-phosphate dikinase and malate dehydrogenase, both associated with the leaf photosynthetic pathway (Fig. 3E), and increased the biomass growth rate by 14% each, followed by cytochrome b5 reductase (24%) and acyl-ACP-hydrolase (24%). MDF showed that, with an ATP concentration below 1.8 mM, the photosynthetic pathway was thermodynamically infeasible (Fig. 3F). Both pyruvate-phosphate dikinase and malate dehydrogenase were combined bottlenecks when the ATP concentration was between 4.7 and 5.7 mM. A similar analysis was performed by varying the concentration of NAD^+^, revealing photosynthetic pathway was thermodynamically feasible when its concentration was between of 0.1 to 5.7 mM. Pyruvate-phosphate dikinase and malate dehydrogenase were the combined bottleneck in this range of NAD^+^ concentration. Thus, the MDF analysis connected metabolic bottlenecks with thermodynamic driving forces. Other bottleneck reactions for heat stress are listed in Supplementary Data 5.

### Parameter tuning of bottleneck enzymes

As the relation between bottleneck reactions and thermodynamics was consistent, we explored different parameters of corresponding enzymes to understand the nature of the bottlenecks. For that, we devised a template-based algorithm, Structure Informed enzyme turnover rate(SI–k_cat_), to calculate *k_cat_* (Supplementary Data 6) of two common enzymes of both stress conditions (cytochrome b5 reductase and acyl-ACP-hydrolase), two enzymes for the heat stress (malate dehydrogenase and pyruvate-phosphate dikinase), and one enzyme for cold stress (coniferyl-aldehyde dehydrogenase). Using predicted *k_cat_* values, we determined the relationship between enzyme concentration (*E*) and saturation (*K*), (equation 12), (Supplementary Fig. 13). *k_cat_* of an enzyme may vary depending on the compartmentalization of the enzyme (all other information such as *v_reaction_* and Δ*G* are also compartment-specific). Therefore, the accuracy of *E* values largely depends on the computational framework of estimating *k_cat_*. *SI – k_cat_* is based on experimental evidence (Supplementary Note) coming from maize or other closely related organisms. Thus, we expect a good accuracy of the calculated *E* values. For all the five enzymes, with lower enzyme saturations (0.1-0.2), higher *k_cat_* significantly reduced the required enzyme concentrations. However, once the enzyme saturation passed 0.6, the effect of different *k_cat_* became lower. Thus, one strategy to improve temperature stress could be to engineer bottleneck enzymes to maintain high saturation by improving its affinity to substrates 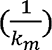.

### Potential of AMF to alleviate bottlenecks: a proof of concept

Although fine-tuning between *k_cat_* and *K* can help improve bottlenecks, it requires significant effort in designing synthetic biology tools. Instead, we can use AMFs, which are well known to alleviate different abiotic stresses in plants^30^. Among these AMFs, the effectiveness of maize inoculation with the *R. irregularis* is well studied^6^. Therefore, as a proof of concept, we generated maize root and leaf transcriptomic data from *R. irregularis* inoculated maize plants to assess the status of bottleneck reactions. In roots, 7769 genes were differentially expressed in inoculated plants compared to control plants, among which 4693 were upregulated (Fig. 4A). Similarly, 6639 genes were differentially expressed in leaves, among which 3485 were upregulated (Fig. 4B). In addition, 1200 and 43 genes showed a log2 fold-change higher than 2 in roots and leaves, respectively. Gene enrichment analysis was then performed for the root genes exhibiting a log2 fold-change higher than 10 (Supplementary Fig. 14). We found an enrichment of xylanase inhibitor protein, suggesting protection from microbial xylanase-induced hemicellulose degradation of maize roots by *R. irregularis*. A similar analysis was conducted with leaf genes exhibiting a log2 fold-change higher than 2 (supplementary fig. 15). We found a significant enrichment of pyruvate dikinase and malate dehydrogenase^31^.

**Fig. 4.**
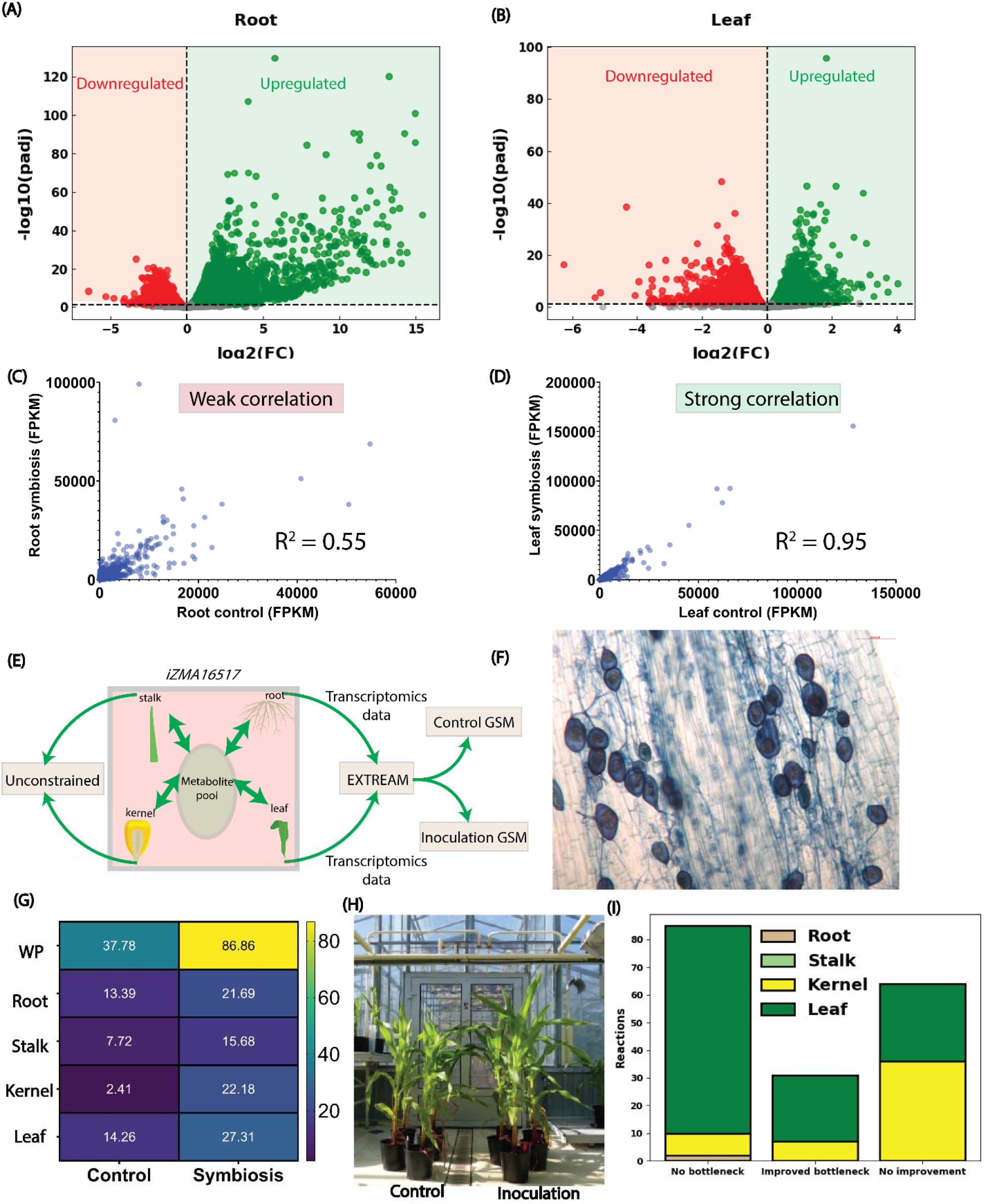
Maize plant responses to the AMF *R. irregularis*. A) Volcano plot of root transcriptomic data for control and inoculated plants. B) Volcano plot of leaf transcriptomic data for control and inoculated plants. C) Comparison between root transcriptomic data for control and inoculated plants. D) Comparison between leaf transcriptomic data for control and symbiotic conditions. E) Root and leaf transcriptomic data integration using the *i*ZMA6517. F) *R. irregularis* inoculation of maize root. G) Root inoculated contextualized iZMA6517 predicted the pattern of biomass growth rate, matched with experimental growth pattern. WP: Whole Plant. H) Picture of plants used for the study displaying higher biomass under inoculation conditions compared to control. I) In the symbiotic interaction, 65% of bottlenecks, identified in cold stress, were alleviated.

We plotted root transcripts for inoculated plants and non-inoculated plants and found a relatively low correlation of 0.55, indicating a distinct transcriptional response after inoculation (Fig. 4C). Alcohol dehydrogenase and pyruvate decarboxylase were significantly upregulated in the symbiosis. For leaves, we found a strong correlation of 0.95, indicating a similar transcriptomic response in the plants inoculated with *R. irregularis* compared to the control (Fig. 4D).

As leaf and root tissues were most affected by the temperature stress, we used these transcriptomics data to reconstruct contextualized *i*ZMA6517s for control and inoculated plants using EXTREAM (Fig. 4E). As these models were not constrained for kernel and stalk, it enabled us to find all theoretical temperature stress alleviation strategies for these two organs. Next, from contextualized models, we found, both whole plant biomass and organ-specific biomass growth rates were higher in the inoculated plants (Fig. 4F-H). Subsequently, implementing MBA to the symbiosis model showed that all the cold stress bottleneck reactions were alleviated. For heat stress, 85 out of 180 (47%) bottleneck reactions were no longer bottlenecks following inoculation (Fig. 4I). Moreover, 31 bottleneck reactions (18%) showed improvement in inoculated plants. Finally, 64 bottleneck reactions (35%) did not show improvement upon inoculation. However, out of those 64 reactions, 62 had a minimal (1-3%) impact on biomass growth rate, except for pyruvate kinase in two different pyruvate 2-O-phosphotransferase of the leaf (8% and 7%), (Supplementary Data 7). Thus, inoculation with *R. irregularis* has the potential to alleviate temperature stress.

## Discussion

Climate change, causing temperature stress, is a leading cause of reduced maize production^32^. Here, we introduced *i*ZMA6517, a multi-organ maize GSM, to better understand the maize metabolism under temperature stress. Fig. 5 shows the overall workflow of this study.

**Fig. 5.**
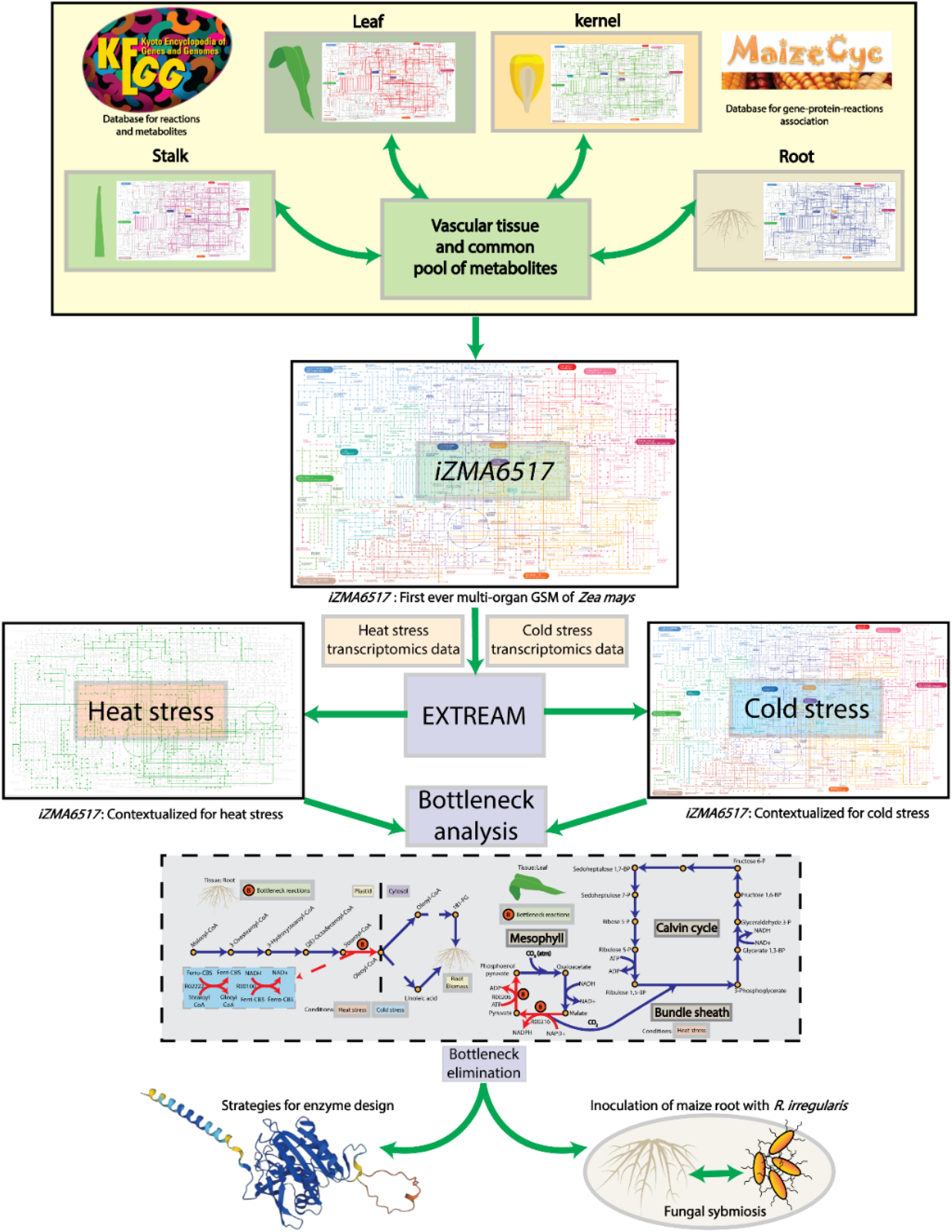
Overall workflow of this project. We reconstructed the first ever multi-organ GSM of maize, *i*ZMA6517. We integrated heat and cold stress transcriptomics data with *i*ZMA6517 with the EXTREAM. Later, we devised MBA to find metabolic bottlenecks of heat and cold stress conditions. We showed that, metabolic bottlenecks on both conditions are guided by thermodynamic principles. We then proposed protein engineering strategies to improve metabolic bottlenecks. Finally, we showed that *R. irregularis* symbiosis with maize root can also alleviate major metabolic bottlenecks.

In first-generation plant GSMs, all organ-specific reactions were combined into a single multi-compartment model. Example of such GSMs are AraGEM^33^, C4GEM^34^, and *i*RS1563^35^. However, these models were unable to simulate inter-organ interactions. The first multi-organ GSM was reconstructed for barley^7^, and subsequently for other plants (see Introduction). However, apart from a core arabidopsis GSM^10^, none of the multi-organ GSMs included root, stalk, kernel, and leaf (Supplementary Data 8). *i*ZMA6517 is the only model thus far combining these four organ-specific GSMs with finer resolution (Supplementary Fig. 16), making it the most comprehensive multi-organ plant GSM.

The E-flux algorithm predicted accurate root phenotype under nitrogen starvation^14^. However, it could not predict the carbohydrate profile in the leaf (Fig. 1F). We hypothesized that the solution space of the E-flux algorithm was overly permissive, resulting in an inaccurate carbohydrate profile. Thus, we further restricted the feasible solution space by equally distributing the transcript of each gene based on the number of reactions the gene participated in (Supplementary Note). The objective function was to minimize the sum of reaction fluxes compared to a reference condition, calculated from the transcriptomic data. A similar objective function was used in MOMA^36^. However, MOMA used wild-type flux distribution as the reference to estimate flux distribution after gene knockouts. In our case, the purpose of the reference condition was to maximize agreement between transcriptomic data and reaction flux. After these modifications, the new algorithm, EXTREAM, predicted the correct carbohydrate profile across leaf cross-sections.

After contextualizing *i*ZMA6517 with EXTREAM, we devised MBA to find plant-wide metabolic bottlenecks. MBA identifies bottleneck reactions in a given metabolic network by expanding the flux space of each reaction to a maximum possible value individually and assess its impact on the biomass growth rate. Compared to the shadow price analysis, which is a metabolite-centric approach, MBA is a reaction-centric approach and better suited for metabolic engineering/bottleneck identification purpose. Under heat stress, 180 reactions were identified as metabolic bottlenecks, whereas only 5 metabolic bottlenecks were identified under cold stress, revealing a fundamental difference between both stresses. This finding can explain the weak transcriptomic correlation between heat and cold stresses (Fig. 1C). MBA indicated that leaf tissue hosted most bottleneck reactions under heat stress (Fig. 3A), consistent with previous works^37^. For cold stress, root tissue hosted most bottleneck reactions (Fig. 3A), also confirmed by an earlier study^38^. To further understand both stresses, we analyzed heat stress bottleneck reactions and 17% of those (Supplementary Data 9) were associated with NAD+/NADH pair, suggesting a relationship between reducing power and heat stress. Additionally, 25% of the bottleneck reactions (Supplementary Data 10) were associated with ADP/ATP pairs. Thus, heat stress is driven by two major components, reducing power and energy generation. Previous studies independently confirmed the effect of reducing power^39^ and energy generation^40^ on heat stress. However, this work showed the synergistic impact of both metabolic components on heat stress. A similar analysis on cold stress bottleneck reactions revealed 60% of those were associated with NAD+/NADH (Supplementary Data 11). Surprisingly, none of the reactions were related to the ADP/ATP pair. Previous studies also confirmed the effect of reducing power on cold stress^41^. We performed the MDF analysis to find common features behind the interplay of reducing power and energy generation. The analysis found all the tested reactions can be the thermodynamic bottlenecks in their respective pathways, revealing a multi-faceted characteristic of temperature stress, geared by the thermodynamics-reducing power-energy generation axis. Literature evidence suggested that cytochrome b5 reductase (one of the two bottleneck enzymes on both stresses) of arabidopsis mutant had lower growth than the wild type^42^. A similar observation was made for acyl-ACP-hydrolase (another common bottleneck enzyme under both heat and cold stress) for arabidopsis^43^. These evidences further validate the proposed thermodynamics-reducing power-energy generation axis.

Enzymes are increasingly repurposed by rational design for directed evolution. Enzyme turnover rate (*k_cat_*) and saturation (*K*) are common parameters for such rational design^44^. A previous work^45^ established a relation among enzyme concentration (*E*), *k_cat_*, and *K*. To explore the relationship for bottleneck enzymes mentioned in the result section, reliable values of *k_cat_* are needed, based on which, *E* can be calculated. There is a deep learning-based algorithm^46^ that can predict *k_cat_*, however, for maize, the algorithm returned *k_cat_* values, not in good agreement with the literature (Supplementary Fig. 17-21). Thus, we proposed a structural similarity weightage-based algorithm (*SI – k_cat_*), which predicted *k_cat_* values of enzymes, close to experimental observations. With the new *k_cat_* values, we sampled the different enzyme saturation to calculate *E*. For all the tested enzymes, the effect of *k_cat_* was prevalent in the lower *K* (Supplementary Fig. 16). However, with the increasing *K*, the effect of *k_cat_* became less. Hence, a strategy for rational enzyme design for alleviating temperature stress can be to implement directed evolution for heat/cold stress to fine-tune the relationship between *k_cat_* – *K*.

A practical approach to alleviating abiotic stress can be to inoculate maize root with an AMF such as *R. irregularis*^30,47–49^. Previous work indicated that AMF inoculated C_4_ plant species exhibited better abiotic stress tolerance through improved quantum yield of PSII, thus providing additional electron sinks^50^. Thus, as a proof of concept, we generated transcriptomic data for control and AMF inoculated maize and reconstructed GSMs for both conditions to assess the status of bottleneck reactions. Inoculation GSM predicted a higher biomass growth rate for all tissues (Fig. 4F). Moreover, we predicted that the inoculation alleviated all cold stress and 65% heat stress reactions. We also calculated flux sum, a proxy for metabolite concentration, for NAD+ and found 49% (Supplementary Fig. 22) increase in the inoculation condition, indicating an additional availability of electron sink provided by the inoculation with *R. irregularis*, which can potentially help maize overcoming the temperature stress. However, *R. irregularis* symbiosis still had bottlenecks from purine, pyruvate, pyrimidine, folate, and fatty acid metabolism (Supplementary Data 12, Supplementary Fig. 23). Additional improvements of the plant biomass production can be achieved by debottlenecking these reactions.

Overall, the first muti-organ maize GSM, *i*ZMA6517, with the aid of EXTREAM and MBA, dissected the impact of temperature stress on maize. Our analysis revealed three major conclusions: (1) Heat and cold stresses are fundamentally different; (2) Both stresses are associated with reducing power, while heat stress has additional bottlenecks in energy generation; and (3) Inoculation with *R. irregularis* can be an effective way to alleviate temperature stress. Using these inferences, a better temperature stress-tolerant maize ideotype with an improved grain yield can be designed. Future work can be extended to elucidate the plant-wide impact of different other abiotic stresses and how abiotic stresses can be ameliorated with the inoculation of AMF. One such case can be to assess the plant-wide impact of *R. irregularis* under high and low nitrogen conditions. This work is currently underway with promising predictions from *i*ZMA6517.

## METHODS

### K-means clustering analysis

K-means clustering^51^ algorithm was used to classify different genes into different clusters. Number of clusters was determined using the Elbow method. The whole K-mean clustering was implemented in Python, using numpy, pandas, and sklearn modules. Default setting of K-mean clustering, mentioned in the sklearn, was not changed in this study. Number of clusters were determined using the elbow method (supplementary Fig. 24)

### GSM reconstruction

A previously published leaf genome-scale model^22^ was used as a scaffolding to reconstruct the stalk and kernel GSMs. These GSMs were later connected with the previously reconstructed leaf^22^ and root^14^ GSMs through vascular tissue to assemble the *i*ZMA6517. Details can be accessed in the supplementary note.

### EXTREAM algorithm

In this work, we proposed the EXTREAM, where transcript of each gene was equally divided based on the number of reactions the gene participated. We also changed the objective function which is the minimization of sum of reaction fluxes compared to a reference condition, calculated from the transcriptomics data.

The formulation of EXTREAM is the following:

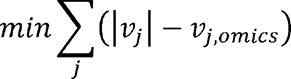

Subject to,

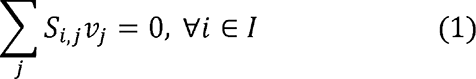

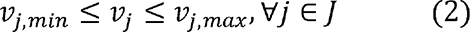

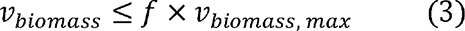

Here, *v_j_* is the flux to be calculated for reaction *J*, *v_j,omics_* is the reference condition calculated from the gene-protein-reaction association for reaction *J*, *S_i,j_* is the stoichiometric matrix for metabolite *I*, and reaction *J*, *v_j,min_* and *v_j,max_* are the upped and lower bound of reaction *J*, *v_biomass_* is the desired biomass growth rate, *v_biomass,max_* is the maximum possible biomass growth rate, and *f* is fraction between 0 to 1. Supplementary note provides the linear reformulation of EXTREAM.

### Metabolic bottleneck analysis

To determine the metabolic bottleneck in a GSM, we proposed the following algorithm.

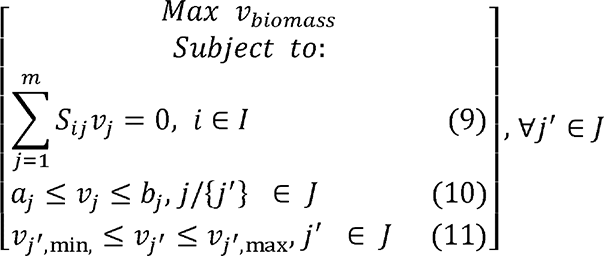

Here *a_j_* is the lower bound reaction *v_j_* and *b_j_* is the upper bound of reaction *v_j_*. Both *a_j_* and *b_j_* were calculated from the transcriptomics data and gene-protein-reaction association. *v_j′,min_* is the expanded lower bound of the reaction *j′* and *v_j′,max_* is the expanded upper bound of the reaction *j′*. In this case, we set 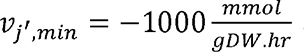 and 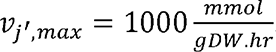. We solved the optimization problem by maximizing the biomass *v_biomass_* for the new expanded flux space of each reaction *j′* in an iterative manner and then recorded the biomass growth rate. From this biomass growth rate collections, we can check for which *j′* biomass growth rate increased significantly. Then that *j′* can be considered as the metabolic bottleneck of a given metabolic network.

### Structure informed *k_cat_* prediction (SI-*k_cat_*)

We calculated the *k_cat_* through scrapping experimental *k_cat_* values from SABIO-RK^52^, structural modeling of enzyme through RGN2^53^, tertiary structure collection of experimentally resolved enzymes, and structural similarity weightage calculation. The protocol can be accessed in the supplementary note along with the validation (supplementary fig. 17-21). The following equation^45^ was used to determine the relationship between enzyme concentration (*E*) and saturation (*K*):

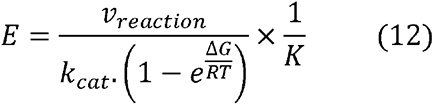

Here, Δ*G* values were collected from the corresponding MDF analysis, and *v_reaction_* were collected by solving contextualized *i*ZMA6517 for heat and/or cold stress.

### *R. irregularis* symbiosis of maize root

The maize line used in this study is B73. The fungal strain used is the highest quality of purity strain *Rhizophagus irregularis* DAOM197198 (Agronutrition, Montpellier, France). In all experiments, maize seeds were surface-sterilized as follows. Seeds were first incubated in ethanol for 5 minutes at 28°C followed by rinsing with distilled water. Then the seeds were incubated for 45 minutes in a 15% commercial bleach solution with 0.01% Triton x100 then rinsed in distilled water. They were placed on wet sterile Watman paper in Petri dishes closed with parafilm and incubated in the dark for 48-72h at 20°C. Germinated seeds were selected and then sown in the experimental setup. The experimental design and the different steps of growing and harvesting for RNAseq experiment, performed in the year of 2020, are indicated in Supplementary Fig. 25. Further details on plant growth conditions and inoculation, harvesting procedures, RNA extraction for RNAseq, bioinformatic analysis, and raw data cleaning can be accessed in the supplementary notes.

## DATA AVAILABILTY

*i*ZMA6517, EXTREAM, and MBA codes are available in this GitHub directory: https://github.com/ssbio/iZMA6517. SI-*k_cat_* codes are available in this GitHub directory: https://github.com/ChowdhuryRatul/kcat_iZMA6517. RNAseq project is deposited in Gene Expression Omnibus (GSE235654). All steps of the experiment, from growth conditions to bioinformatic analyses, were detailed in CATdb: http://tools.ips2.u-psud.fr.fr/CATdb/; Project: NGS2021_19_Rhizophagus according to the MINSEQE. Nutrients for the plant growth can be accessed in Supplementary Data 13.

## ACKNOWLEDHEMENT

RS gratefully acknowledges funding support from National Science Foundation (NSF) CAREER grant (1943310) and Nebraska Collaboration Initiative Grant (21-1106-6011). CDM acknowledges partial support by the Center for Bioenergy Innovation, a U.S. Department of Energy Bioenergy Research Center supported by the Office of Biological and Environmental Research in the DOE Office of Science. RC is grateful for Iowa State University startup grant and Translation AI Center (TrAC) Seed grant. AD acknowledges the support from Plant2Pro® by the French National Agency for Research - ANR (agreement #18-CARN-024-01 – 2018). AD also acknowledges the benefits received from France 2030 program (Saclay Plant Sciences, reference n° ANR-17-EUR-0007, EUR SPS-GSR integrated into France 2030 (reference n° ANR-11-IDEX-0003-02), INRAE Grant IB BAP 2021 MYCORN, and IJPB’s Plant Observatory technological platforms. We acknowledge the constructive comments of Margaret Simons-Senftle during the model reconstruction process. We thank Michel Lebrusque, Lilan Diahuron, Patrick Grillot, and Christian Jeudy for taking care of the plants in the greenhouse. We thank the POPS platform for the RNAseq experiment and data. We also thank Etienne Delannoy, Christine Paysant le Roux, and Alexandra Launay-Avon from POPS platform for fruitful discussions. We are grateful to Cyril Bauland and Carine Palaffre from the INRAE for providing B73 seeds from the Centre de Ressources Biologiques of Saint Martin de Hinx (France). The original B73 seeds were provided by the North Central Regional Plant Introduction Station (IA, USA) for whom we are grateful.

## AUTHOR CONTRIBUTION

R.S., C.D.M., A.D., R.C., and B.H. conceived the study and edited the manuscript. N.B.C. wrote the manuscript. N.B.C reconstructed and curated the model. K.A.S. and B.A. generated the *k_cat_* values. N.B.C performed all computational analyses. B.D. performed the experiments on the inoculation study. I.Q. was involved in experimental design and plant care. M.R. was involved in plant growth and care, and monitoring RNA extractions.

## ADDITIONAL INFORMATION

Supplementary information is available for this paper at the online version.

